# AbNovoBench: a resource and benchmarking platform for monoclonal antibody de novo sequencing

**DOI:** 10.64898/2026.02.02.703105

**Authors:** Wenbin Jiang, Ling Luo, Yueting Xiong, Jin Xiao, Zihan Lin, Lihong Huang, Yiie Qiu, Sainan Zhang, Jingyi Wang, Chao Wang, Ningshao Xia, Quan Yuan, Rongshan Yu

**Author notes:** These authors contributed equally to this work (Wenbin Jiang, Ling Luo, Yueting Xiong, Jin Xiao).

## Abstract

Monoclonal antibodies (mAbs) are critical in disease diagnostics and therapeutics, yet the performance of mass spectrometry (MS)-based de novo sequencing remains incompletely characterized due to limited antibody-specific datasets and the absence of a standardized benchmark framework. Here we present AbNovoBench, a comprehensive framework for evaluating data analysis strategies for mAb de novo sequencing. It features the largest high-quality dataset to date, generated in-house, comprising 1,638,248 peptide-spectrum matches from 131 mAbs across six species and 11 proteases, supplemented by eight mAbs with known full-length sequence for end-to-end reconstruction assessment. Employing a unified training dataset, we systematically benchmarked 13 deep learning-based de novo peptide sequencing algorithms and three assembly strategies across peptide sequencing metrics (accuracy, robustness, efficiency, error types) and assembly metrics (coverage depth, assembly score). AbNovoBench (https://abnovobench.com) provides an online platform enriched with curated antibody MS resources and pre-trained models, enabling customizable antibody sequencing workflows, accelerating antibody-specific algorithms development, and improving reproducibility in proteomics.

## 1 Introduction

Monoclonal antibodies (mAbs) are engineered immunoglobulins designed to recognize defined antigenic epitopes, thereby recapitulating key features of natural immune responses[1]. Owing to their high specificity, affinity, and experimental reproducibility, mAbs have become indispensable across basic research, diagnostics applications, and therapeutic development[2, 3]. However, insufficient sequence confirmation and quality control continue to undermine research reproducibility and compromising the reliability of antibody-based discoveries[4, 5]. Antigen-binding versatility primarily arises from the complementarity-determining regions (CDRs) within the variable domains[6], making accurate antibody sequence information fundamental to understanding structure–function relationships, immune diversity[7], and antibody engineering strategies such as cross-species constant-region exchange to modulate specificity and expand functional diversity.

Despite their central importance, obtaining complete and accurate monoclonal antibody sequences remains challenging. Traditional approaches depend on mRNA derived from hybridomas[8] and therefore fail to capture the protein-level heterogeneity of antibodies, neglecting critical post-translational modifications (PTMs) that influence binding affinity, developability, and effector functions[9]. Mass spectrometry (MS)-based de novo sequencing of secreted antibodies provides a powerful complementary strategy that overcomes these limitations[10, 11]. Bottom-up proteomics workflows enzymatically digest antibodies into peptides, acquire tandem mass spectra by LC-MS/MS, and reconstruct full-length heavy and light chains through reference-free sequencing and assembly, enabling antibody recovery even from lost or unavailable hybridomas and laying the foundation for large-scale serological sequencing.

Since the introduction of DeepNovo[12] in 2017, deep learning (DL) has transformed de novo peptide sequencing by learning direct mappings from MS/MS spectra to amino-acid sequences. The pioneering model integrated convolutional and recurrent networks for effective encoding and prediction[12], while SMSNet[13] bridging de novo and database methods via confidence-based refinements. Subsequent innovations include: order-invariant processing of m/z-intensity pairs (PointNovo[14]); transformer self-attention with precursor embeddings (Casanovo[15, 16]); parallelized point cloud architectures for efficiency (PGPointNovo[17]); diffusion models (InstaNovo[18]); convolutional architectures (PepNet[19]); amino acid-aware transformers (DPST[20]); graph networks for ion gaps (GraphNovo[21]); spectral regularity detection (PowerNovo[22]); mass-sensitive convolutions (Spectralis[23]); complementary ion reconstruction (*π*-HelixNovo[24]); bidirectional decoding (NovoB[25]); adaptive PTM training (AdaNovo[26]); contrastive learning (ContraNovo[27]); and non-autoregressive, accelerated transformers (*π*-PrimeNovo[28]). Nevertheless, peptide-level identification does not directly translate into accurate protein-level assembly. Although antibody-specific assembly frameworks such as ALPS[29], Stitch[30, 31], and Fusion[32] have partially alleviated this challenge by integrating de novo reads with germline templates, critical issues remain unresolved, including full-length sequence fidelity and reliable reconstruction of highly variable regions, particularly CDR3.

A comprehensive and standardized benchmarking framework is therefore urgently needed to systematically evaluate de novo peptide sequencing methods in the context of monoclonal antibody reconstruction. Currently DL-based peptide sequencing algorithms are not specifically designed for antibody data, are often trained on heterogeneous and non-standardized datasets, and are typically evaluated using limited metrics focused on peptide- or amino-acid-level accuracy. For instance, key properties relevant to antibody sequencing, such as PTM identification, coverage across variable domains, and computational efficiency, remain insufficiently assessed, often on datasets comprising only a few antibodies. To address these gaps, we developed AbNovoBench, a systematic benchmarking framework tailored for mAb de novo sequencing. By integrating a streamlined single-pot multi-enzymatic gradient digestion (SP-MEGD) strategy[32] with multi-species, multi-protease designs, AbNovoBench generates diverse, high-quality antibody MS/MS datasets and introduces antibody-specific evaluation metrics. Together, this framework establishes standardized datasets and protocols that enable rigorous comparison of existing tools and facilitate the development of next-generation antibody-aware sequencing algorithms.

## 2 Results

### 2.1 An antibody-specific dataset for mAb de novo sequencing evaluation

Antibody-specific datasets are essential for the rigorous evaluation of mass spectrometry-based de novo sequencing methods for mAbs. To this end, we constructed a dedicated benchmarking dataset utilizing the previously developed single-pot multi-enzymatic gradient digestion (SP-MEGD) protocol [32]. This approach integrates multiple proteases within a single-vessel, time-resolved reaction, enabling efficient generation of dense and informative antibody peptide coverage. Combined with multi-species and multi-protease experimental design, the resulting dataset comprises 1,638,248 peptide-spectrum matches (PSMs) derived from 131 mAbs, spanning 6 species and 11 proteases (Fig. 1 and Fig. S1). Additionally, we generated complete MS/MS datasets for eight mAbs with known full-length sequences, enabling systematic evaluation of integrated peptide sequencing and assembly strategies in terms of reconstruction completeness and accuracy (Table S1).

**Fig. 1.**
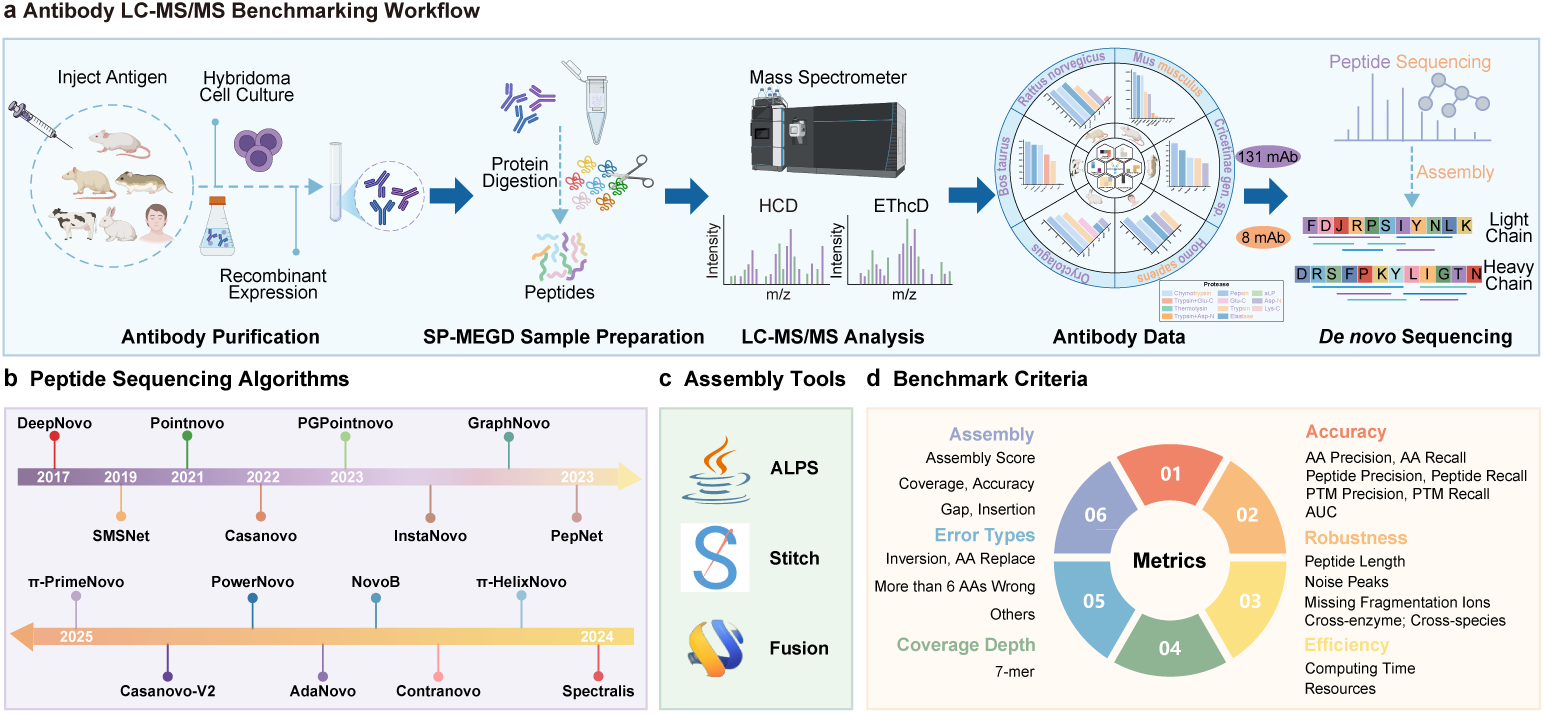
Overview monoclonal antibody de novo sequencing analysis benchmark. **a.** Schematic of the antibody LC-MS/MS benchmarking workflow. Monoclonal antibodies were generated via hybridoma or recombinant expression methods, purified, and digested using various proteases with SP-MEGD method. Purified peptides were identified using a dual fragmentation strategy (HCD & EThcD). Two benchmark datasets were established: a PSM-level dataset comprising 131 antibodies with known peptide sequences and diverse proteases, and a full-spectrum dataset of 8 antibodies for downstream assembly evaluation. **b.** Deep learning methods for de novo peptide sequencing from 2017 to 2025. **c.** Assembly tools evaluated in this benchmark: ALPS (fully de novo), Stitch and Fusion (both template-based). **d.** Benchmarking criteria spanning accuracy (including AA-, peptide-, PTM-level metrics, and AUC), robustness (tolerance to different influencing factors), efficiency (runtime and resource usage), error types, coverage depth, and final assembly quality.

Compared with a recent antibody-focused benchmarking study [3], our dataset exhibits substantially greater scale and spectral quality. That study evaluated peptide sequencing and assembly using only 25,960 PSMs obtained from three publicly available monoclonal antibodies, with some experiments relying on separately acquired heavy and light chains—an experimental design that deviates from current practical workflows. Moreover, the underlying data were generated using earlier-generation instrumentation, resulting in limited spectral quality: 90.51% of validated spectra lacked at least one fragment ion, and 84.32% of peaks were classified as noise. By contrast, in our dataset, only 44.08% of spectra exhibit missing fragment ions, and the proportion of noise peaks is reduced to 76.56% under the same criteria, reflecting a marked improvement in data quality. Together, this high-quality, standardized antibody-specific dataset provides a robust foundation for systematic and unbiased evaluation of de novo sequencing and assembly methods. Beyond enabling rigorous performance comparisons among existing tools, it facilitates the development of more accurate and generalizable algorithms tailored to the unique complexity of antibody proteomics.

### 2.2 Accuracy metrics for antibody PSM de novo sequencing

To enable standardized evaluation, we retrained a panel of published de novo peptide sequencing algorithms using a uniformly filtered dataset derived from the MassIVE-KB-v130 database. As summarized in Table S2, several models exhibited unstable training behaviour and were therefore excluded from further analysis. The final benchmark included 13 algorithms, spanning diverse architectural paradigms, including recurrent neural network–based, transformer-based, and point cloud-based frameworks. Specifically, the evaluated methods comprised DeepNovo [12], SMSNet [13], Casanovo (v1 [15] and v2 [16]), ContraNovo [27], InstaNovo [18], PepNet [19],

AdaNovo [26], PointNovo [14], PGPointNovo [17], pNovo3 [33], *π*-HelixNovo [24], and *π*-PrimeNovo [28]. Notably, for ContraNovo [27] and *π*-PrimeNovo [28], we also included the authors’ original models trained on the full MassIVE-KB-v130 dataset, as retraining on the refined subset resulted in degraded performance.

We first evaluated each algorithm using seven complementary accuracy metrics at different granularities: amino acid (AA) precision and recall, peptide-level precision and recall, post-translational modification (PTM) precision and recall, and the area under the precision–recall curve (AUC), which captures overall de novo sequencing performance (Fig. 2A). Among all evaluated models, ContraNovo achieved the highest scores across multiple categories, including AA precision (0.7843) and recall (0.7860), peptide-level precision and recall (both 0.6696), PTM recall (0.6528), and AUC (0.6106). Interestingly, Casanovo v1, trained on a filtered subset of the MassIVE-KB-v130 dataset (1,913,865 PSMs) containing only three common modifications—oxidation (M) and deamidation (N/Q)—achieved peptide sequencing performance comparable to ContraNovo, despite the latter being trained on the full MassIVE-KB-v1 dataset (30,504,897 PSMs) with a substantially broader modification space. In fact, Casanovo v1 exhibited slightly higher PTM precision (0.7414 vs. 0.6528) and competitive AUC (0.5564 vs. 0.6106) than ContraNovo, suggesting that expanding the training dataset and modification space does not always lead to improved down-stream performance. These findings indicate that a more focused training strategy, centered on biologically relevant PTMs and curated spectra, may be sufficient—and even advantageous—for applications such as monoclonal antibody sequencing, where the chemical landscape is relatively well-defined. Casanovo v2 and InstaNovo also demonstrated robust performance, with Casanovo v2 achieving strong PTM precision (0.7462) and AA recall (0.7311), while InstaNovo exhibited the highest PTM precision across all models (0.7626) along with solid peptide recall (0.6453). In contrast, models such as DeepNovo, PepNet, and pNovo3 underperformed across several metrics, particularly in PTM recall and AUC, reflecting limited adaptability to antibody-specific spectra. SMSNet and AdaNovo achieved moderate results, with relatively higher AA recall but lower peptide-level precision, while PointNovo and PGPointNovo showed minimal improvements over their predecessors. The *π*-HelixNovo and *π*-PrimeNovo models displayed acceptable AA and peptide-level accuracy, yet underperformed in terms of AUC values. Collectively, these results highlight substantial performance differences across algorithmic designs and consistently demonstrate the superiority of transformer-based approaches, particularly ContraNovo and the Casanovo variants, for antibody PSM–level de novo sequencing.

**Fig. 2.**
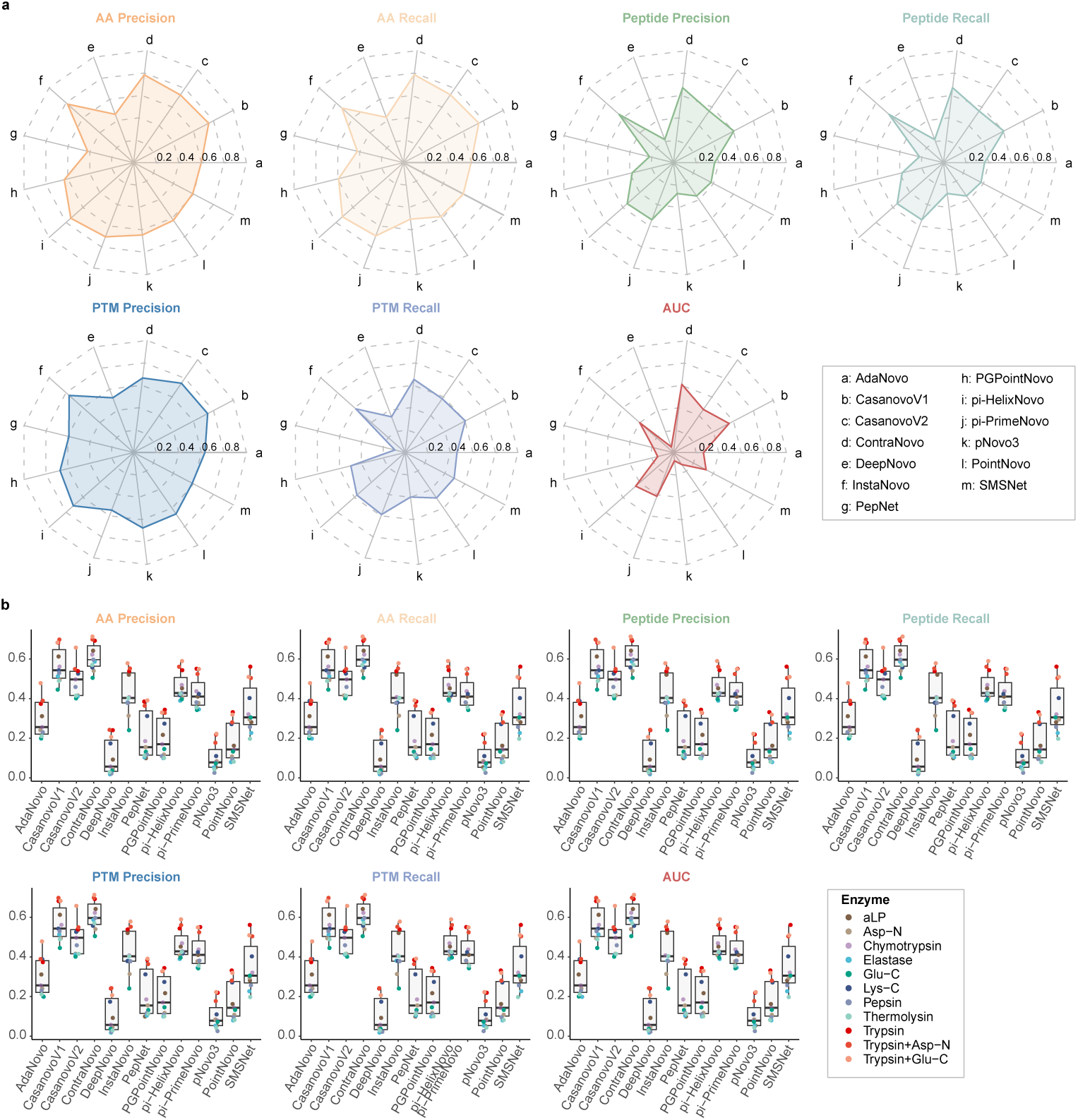
Performance evaluation of de novo peptide sequencing algorithms on monoclonal antibody PSMs. **a.** Performance of 13 algorithms across seven metrics: AA precision, AA recall, peptide precision, peptide recall, PTM precision, PTM recall, and AUC. **b.** Metric variations under 11 different protease digestion strategies.

### 2.3 Evaluation of cross-enzyme and cross-species generalizability

Cross-enzyme robustness is a critical quality attribute of de novo peptide sequencing in bottom-up proteomics. However, most published studies of newly developed sequencing algorithms primarily focus on tryptic datasets, owing to their widespread availability and extensive use in non-antibody proteomics, while largely neglecting performance on non-tryptic proteases. Building upon the accuracy analyses described above, we systematically evaluated enzyme-specific generalization across 11 proteolytic conditions, including three trypsin-based settings (trypsin, trypsin+GluC, trypsin+AspN) and eight non-tryptic proteases (GluC, LysC, chymotrypsin, AspN, thermolysin, elastase, pepsin, and aLP) (Fig. 2B). As expected, models trained predominantly on tryptic spectra achieved their highest accuracy under trypsin digestion. Nevertheless, performance across different proteolytic conditions varied substantially among algorithms. Transformer-based models, including ContraNovo and Casanovo v1/v2, consistently maintained high amino acid- and peptide-level recall across both tryptic and non-tryptic conditions, exhibiting particularly stable performance under challenging cleavage regimes such as thermolysin and AspN. InstaNovo also demonstrated broad robustness, with notably strong PTM recall under GluC, pepsin, and chymotrypsin digestion. In contrast, DeepNovo, PepNet, and pNovo3 showed pronounced performance degradation under several non-tryptic conditions, particularly those involving N-terminal or low-specificity cleavage. PointNovo and PGPointNovo displayed substantial variability across enzymatic contexts, whereas *π*-PrimeNovo, despite being evaluated using a pretrained model, retained moderate recall under GluC and elastase digestion. Collectively, these results reveal marked differences in enzyme-specific robustness across architectural designs and underscore the superior out-of-distribution generalization of transformer-based models.

To further assess biological generalizability, we extended the analysis to antibody PSMs derived from six species, including *Homo sapiens*, *Bos taurus*, *Mus musculus*, *Rattus norvegicus*, *Oryctolagus cuniculus*, and *Cricetinae* (Fig. S2). Although all models were trained exclusively on human-derived spectra, their performance remained largely consistent across species, indicating strong transferability of learned spectral representations. Notably, ContraNovo, Casanovo v1, and InstaNovo maintained high amino-acid– and peptide-level accuracy in both human and non-human datasets. Similarly, PTM precision and recall were stable across species, with only minor variation attributable to species-specific fragmentation patterns or sample preparation differences. This robustness across diverse biological contexts suggests that, when provided with sufficiently high-quality spectra, modern de novo peptide sequencing models can generalize effectively beyond their original training domain.

### 2.4 Biochemical and spectral determinants of de novo peptide sequencing accuracy, and model-specific error patterns

Although intrinsic peptide characteristics and spectral quality are widely recognized as key determinants of de novo sequencing performance, their systematic impact has not been comprehensively quantified. We therefore examined how peptide length, the number of missed cleavages, and spectral noise influence model accuracy. Across all evaluated algorithms, peptide length exhibited a strong negative correlation with peptide-level recall (Fig. 3A), with Pearson correlation coefficients ranging from *−*0.81 to *−*0.98. Performance declined sharply once peptide length exceeded approximately 19 to 20 residues, and for most models, peptide recall approached zero for sequences longer than 25 amino acids. Similarly, an increasing number of missed cleavage sites was strongly associated with reduced recall (Fig. 3B; *R < −*0.85), suggesting that peptide length and incomplete proteolysis independently impair prediction accuracy. The combined effects of these two factors are shown in Fig. S3, where peptides that are both long and contain *≥* 4 missed cleavages consistently yielded recall values below 0.1 across all models. Notably, under these adverse conditions, transformer-based methods, including Casanovo v1/v2, ContraNovo, and InstaNovo, retained comparatively higher recall than other algorithms. We further assessed the impact of spectral quality using a noise factor metric derived directly from the raw MS/MS data (Fig. S4). Peptide recall exhibited a uniformly strong negative correlation with noise level in all models (*R < −*0.96), underscoring the sensitivity of de novo peptide sequencing to signal quality. Collectively, these results identify peptide length, proteolytic completeness, and spectral noise as dominant limiting factors for peptide-level reconstruction, while highlighting the superior robustness of transformer-based models across diverse biochemical and analytical challenges.

**Fig. 3.**
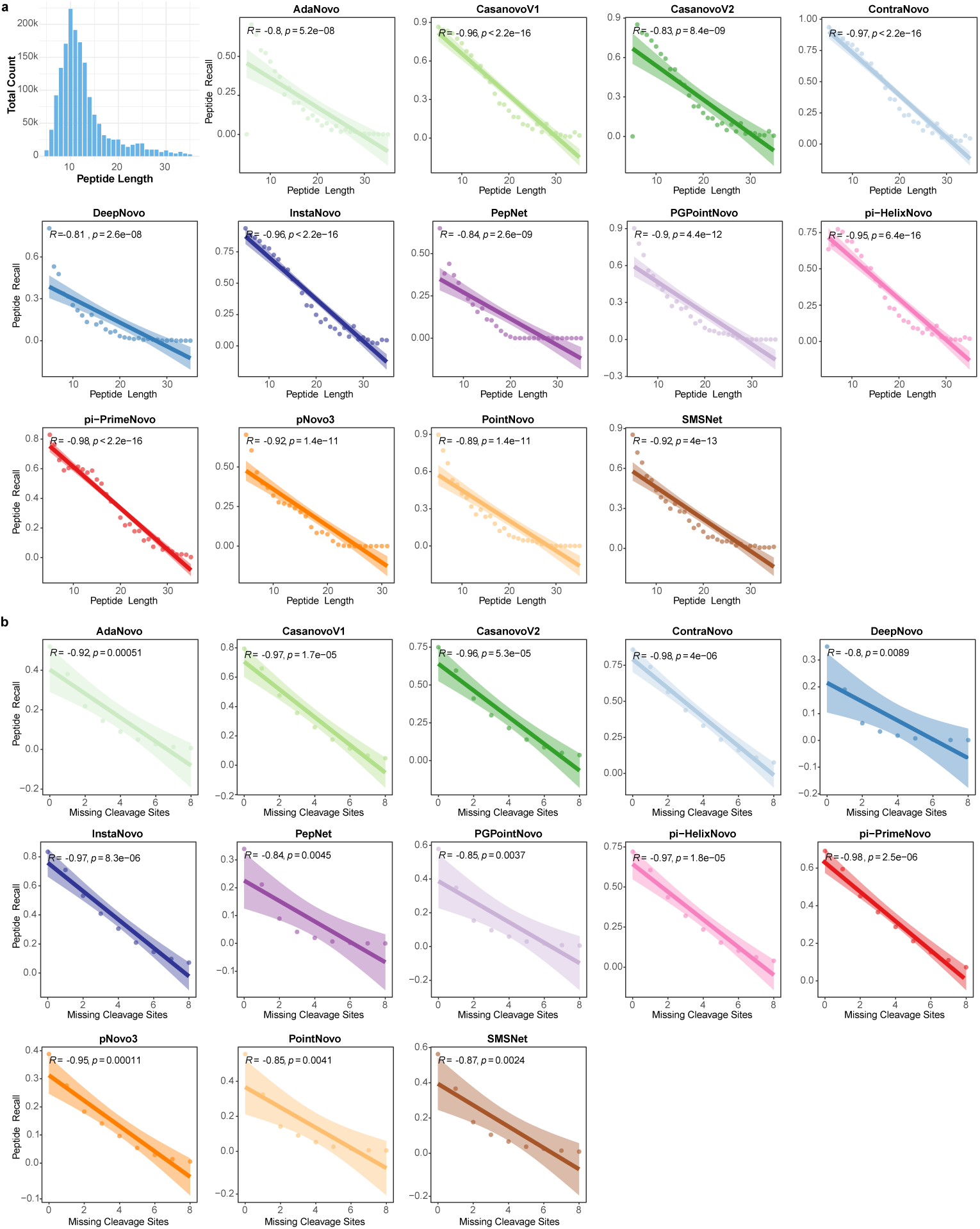
Impact of peptide properties on de novo sequencing performance. **a.** Relationship between peptide length and peptide recall across 13 algorithms. Each dot represents recall for peptides of a given length; shaded areas indicate 95% confidence intervals. **b.** Relationship between the number of missed cleavage sites and peptide recall. Strong negative correlations are observed for both factors across all models.

Beyond aggregate accuracy, we next analysed the sources and structural patterns of incorrect predictions. To systematically characterize failure modes, erroneous peptide predictions were categorized into 11 distinct structural error types, and their frequencies were compared across all antibody PSMs for the 13 evaluated algorithms (Table S3). Across most models, the predominant failure mode was large-fragment misassignment, defined as incorrect prediction of more than six amino acids. This error class constituted the majority of incorrect predictions in DeepNovo (57.72%) and remained highly prevalent in SMSNet (45.80%), PepNet (42.83%), PointNovo (40.83%), and PGPointNovo (40.60%), indicating a systematic tendency toward large-fragment mismatches. Models including AdaNovo, *π*-HelixNovo, Casanovo v1, Casanovo v2, ContraNovo, and *π*-PrimeNovo also exhibited moderate proportions of large-fragment errors (33.76%–39.37%). In contrast, InstaNovo showed a comparatively lower rate of large-fragment errors (31.24%) but displayed a clear bias toward short-range structural disruptions, with inversion of the first three residues as the most frequent error (8.10%), followed by 2-AA (7.38%) and 1-AA (7.20%) substitutions. Although pNovo3 exhibited the lowest proportion of large-fragment errors (28.69%), it showed the highest overall error rate (89.26%) and elevated frequencies across nearly all error categories, reflecting broadly distributed but pervasive inaccuracies. Together, these analyses distinguish models dominated by large-fragment mismatches from those characterized by localized or structurally diverse prediction failures, providing mechanistic insight into the distinct error landscapes underlying de novo peptide sequencing algorithms.

### 2.5 Assembly performance of de novo and template-guided strategies

Accurate full-length sequence assembly remains a major challenge in mAb de novo sequencing. To improve the reliability of downstream antibody sequence assembly, we first evaluated the coverage depth of predicted peptides along the heavy chain, with particular emphasis on the CDRs, which are especially difficult to reconstruct. Because current assembly strategies primarily rely on peptide overlap information, insufficient coverage within CDRs directly disrupts assembly continuity. Using a sliding window of seven residues, we calculated position-specific coverage depth for 13 de novo peptide sequencing algorithms across eight monoclonal antibodies (Fig. S5). To ensure robust assembly input, we required a minimum coverage depth of 10 at all CDR positions; algorithms failing to meet this threshold at any CDR position were deemed insufficiently representative and excluded from further analysis. Under this criterion, five algorithms were removed, leaving eight models—Casanovo v1, Casanovo v2, ContraNovo, InstaNovo, PGPointNovo, *π*-HelixNovo, *π*-PrimeNovo, and PointNovo—for subsequent antibody assembly evaluation.

We first evaluated ALPS [29], a fully de novo assembly approach that reconstructs antibody sequences directly from overlapping peptides without reference templates. As summarized in Supplementary Data Assembly-result ALPS.xlsx, ALPS-generated contigs showed high coverage and accuracy when aligned post hoc to known antibody sequences. However, such alignment-based evaluation implicitly assumes access to the ground-truth sequence, which is typically unavailable in real discovery settings. In the absence of prior knowledge, determining the precise genomic location and correctness of each contig becomes challenging. Even when contigs overlap or align to the same region, distinguishing accurate assemblies from erroneous ones remains difficult. Moreover, analysis of the longest contigs revealed that they rarely achieved complete and accurate coverage of entire antibody chains. Instead, contigs were often confined to constant regions and, in some cases, spanned fragments from both heavy and light chains. These observations indicate that, given the current state of de novo peptide sequencing, fully de novo assembly remains substantially limited for antibodies with unknown sequences.

Given these limitations, we next examined two template-based assembly strategies, Fusion [32] and Stitch [31]. Compared with fully de novo assembly, both algorithms exhibited marked improvements in accuracy and continuity, particularly within the variable CDRs (Fig. 4). Fusion leverages germline templates to resolve ambiguous overlaps and bridge poorly supported regions, yielding fully contiguous chain assemblies without insertions or gaps. In comparison, Stitch achieved comparable chain-level coverage but adopted a more conservative strategy, introducing occasional gaps in regions with limited peptide support. These differences are further quantified by insertion and gap analyses (Fig. S6) and by assembly scores at both whole-antibody and CDR levels (Fig. S7), with the largest advantage observed in CDR3. To assess practical applicability, we further analyzed inference throughput across models (Fig. S8).

**Fig. 4.**
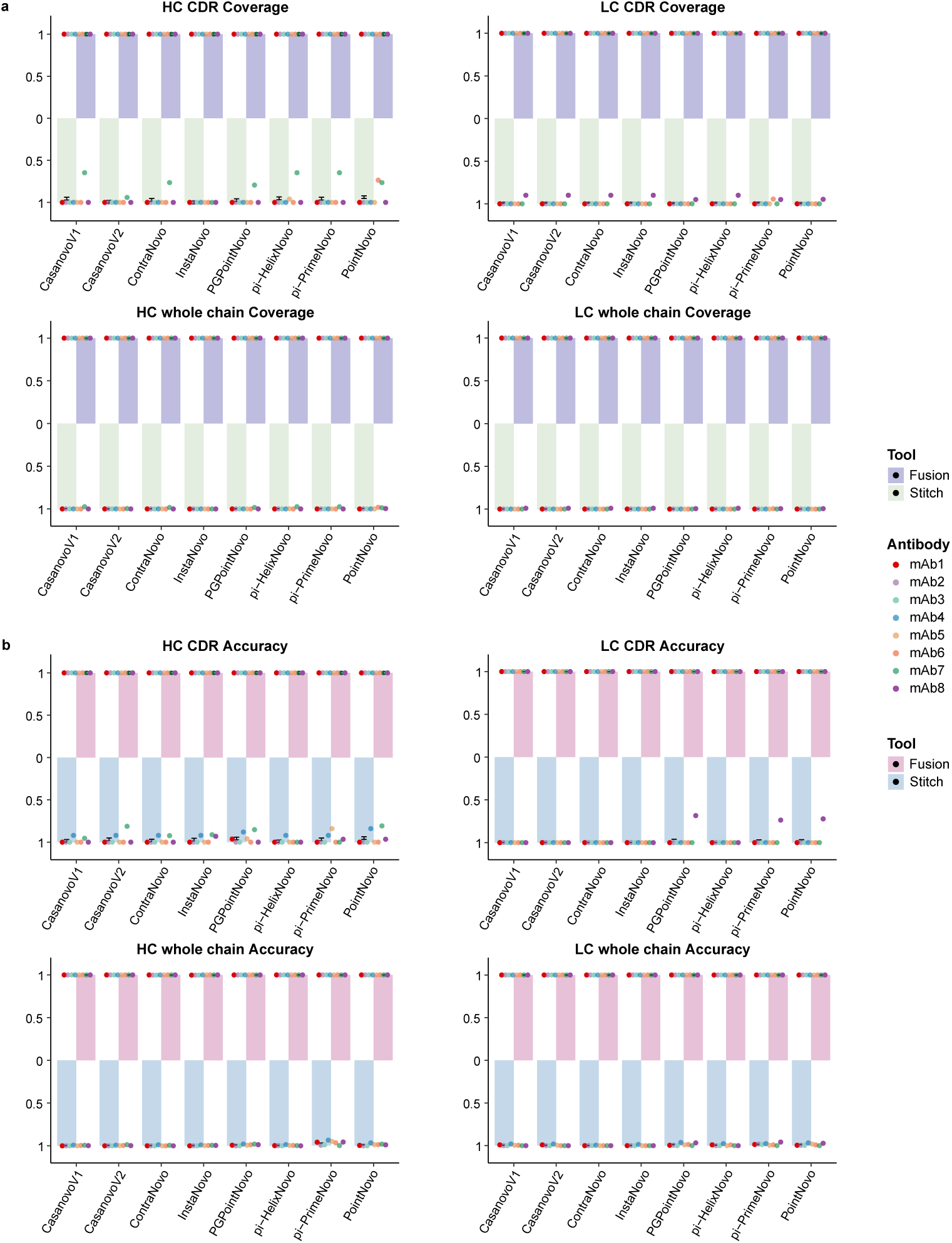
Assembly performance for Stitch and Fusion across eight monoclonal antibodies. **a.** Coverage and **b.** accuracy of HC and LC CDRs, and full-length chains, using the Stitch and Fusion assembly methods.

### 2.6 Development of a benchmarking platform for antibody de novo sequencing

To facilitate comprehensive and standardized evaluation of antibody de novo sequencing algorithms, we constructed two benchmark datasets: a PSM-level dataset for assessing peptide-level accuracy, and a complete MS/MS dataset comprising eight monoclonal antibodies with known full-length sequences for evaluating downstream assembly performance. Building upon these datasets, we developed AbNovoBench (Fig. 5), a web-based benchmarking platform that provides a unified evaluation pipeline, interactive result visualization, and continuous community-driven updates. AbNovoBench supports the submission of newly developed de novo sequencing models as well as retrained versions of existing algorithms, facilitating real-time tracking of methodological advances. Notably, all tools evaluated in this study were bench-marked using their default parameter settings to ensure fair comparison; however, the platform also allows re-evaluation of models optimized with alternative configurations. Additionally, users can directly select appropriate pre-trained models from AbNovoBench to construct customized antibody sequencing workflows. This capability accelerates pipeline development by eliminating the need for model retraining from scratch, thereby reducing both computational cost and development time while leveraging high-quality, pre-optimized models. In summary, AbNovoBench constitutes a scalable and transparent resource for benchmarking antibody de novo sequencing tools. By standardizing evaluation protocols and facilitating open comparison, the platform not only enables rigorous assessment of existing methods but also fosters continued innovation in antibody-specific sequencing algorithms.

**Fig. 5.**
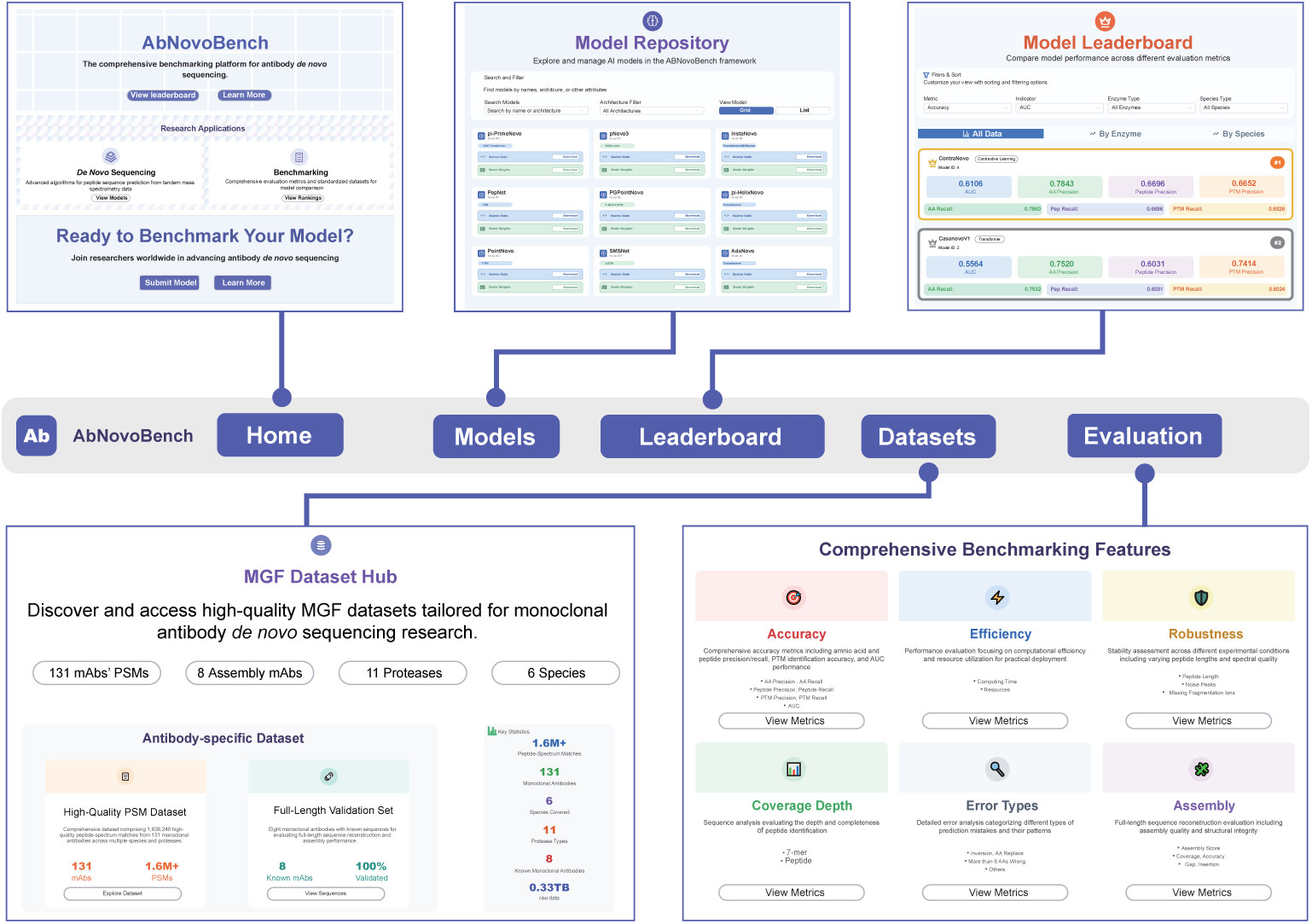
AbNovoBench: a benchmarking platform for monoclonal antibody de novo sequencing analysis. Snapshots of the AbNovoBench web interface illustrating its core functionalities. The interface features a Home page that introduces the platform and provides quick access to the dataset hub, evaluation modules, a Model Repository for browsing available methods and associated model metadata, and a Model Leaderboard that summarizes comparative performance across various benchmark configurations. The MGF Dataset Hub curates downloadable MGF datasets encompassing multiple antibody panels, protease digestion protocols, and species. Employing a unified training dataset, we benchmarked different deep learning-based de novo peptide sequencing algorithms and various assembly strategies across six metric categories, spanning peptide sequencing (accuracy, robustness, efficiency, error types) and assembly (coverage depth, assembly score).

**Fig. S1.**
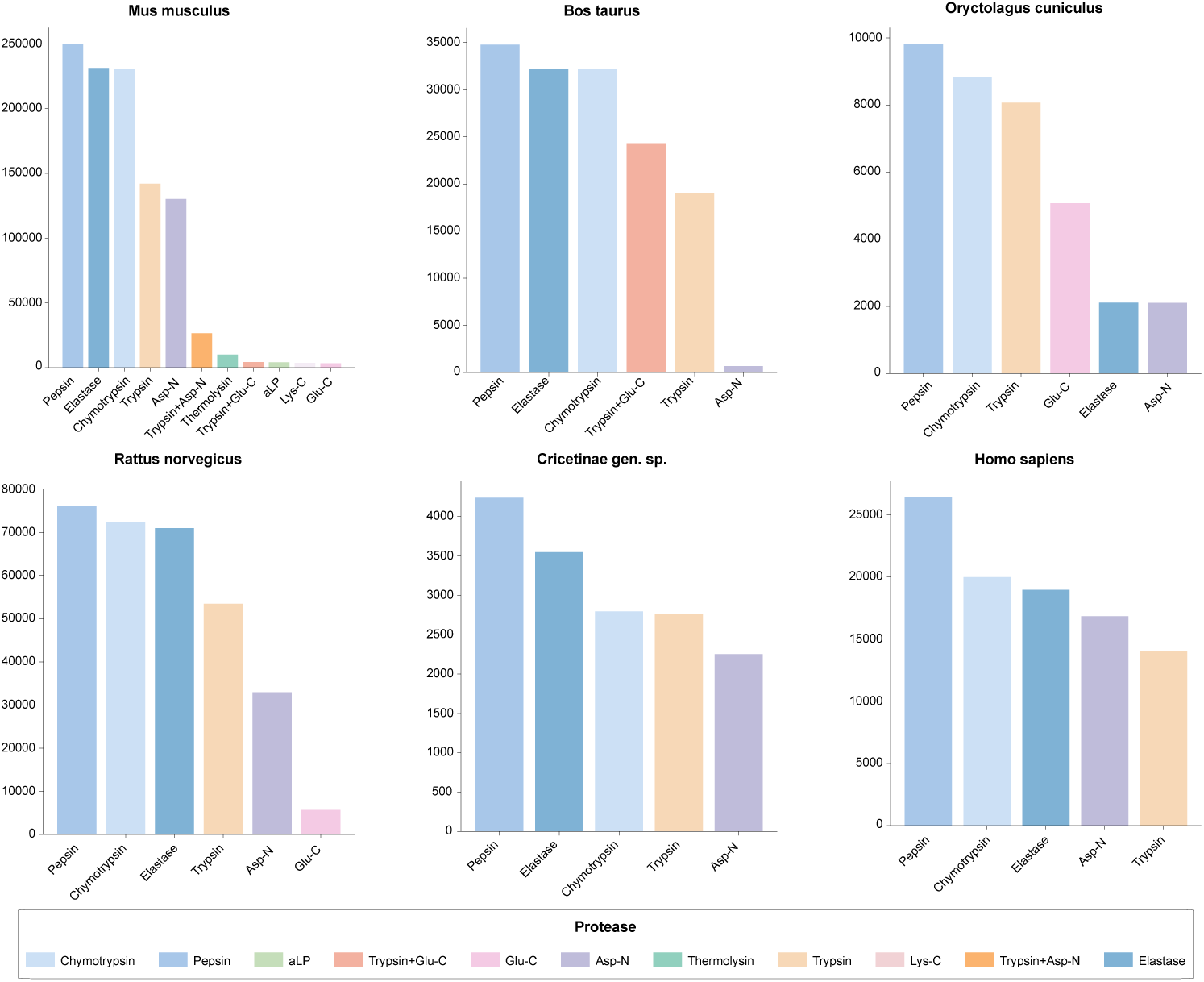
Number of spectra for different proteases across species. Bar plots showing the number of spectra for different proteases across species. Proteases are color-coded according to the legend.

**Fig. S2.**
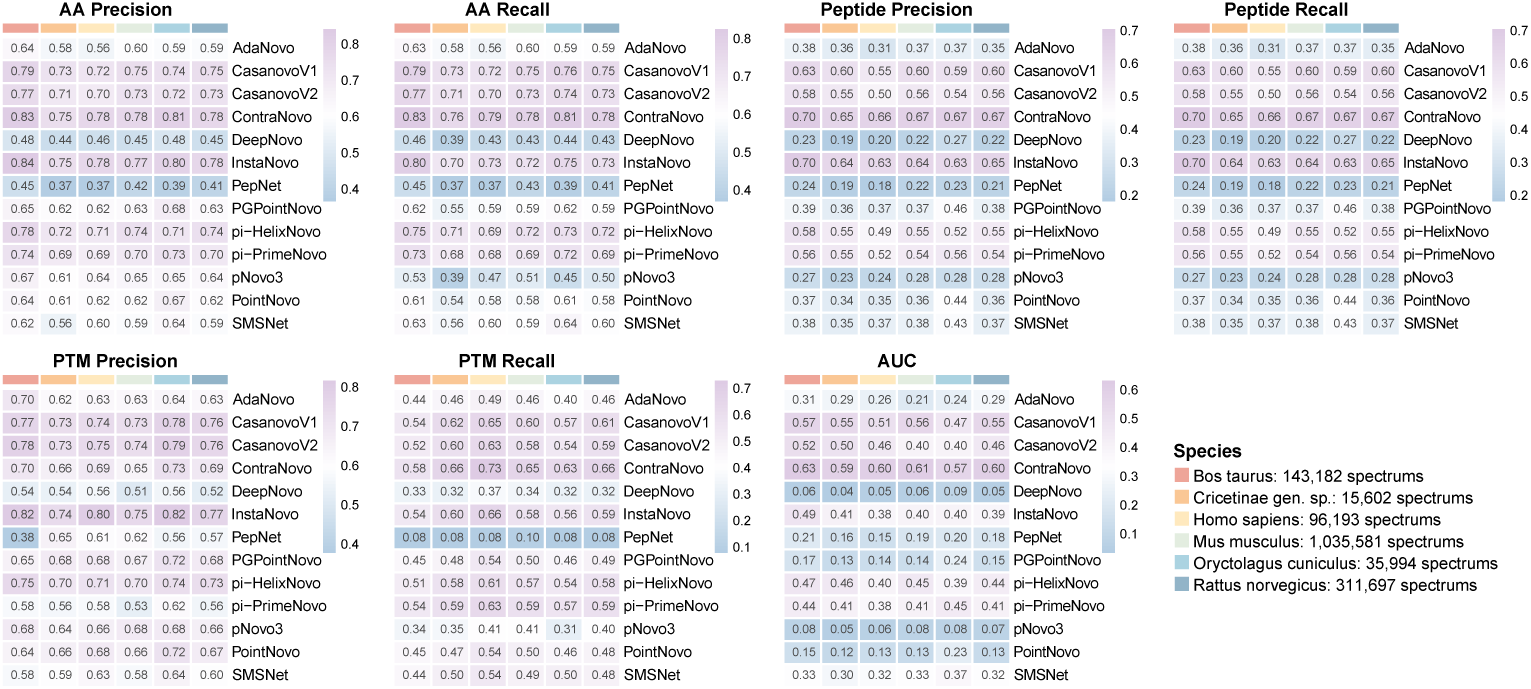
Performance of de novo peptide sequencing algorithms across species. Heatmaps of seven accuracy-related metrics (AA precision/recall, Peptide precision/recall, PTM precision/recall, and AUC) for 13 algorithms across six species. Color intensity indicates metric values.

**Fig. S3.**
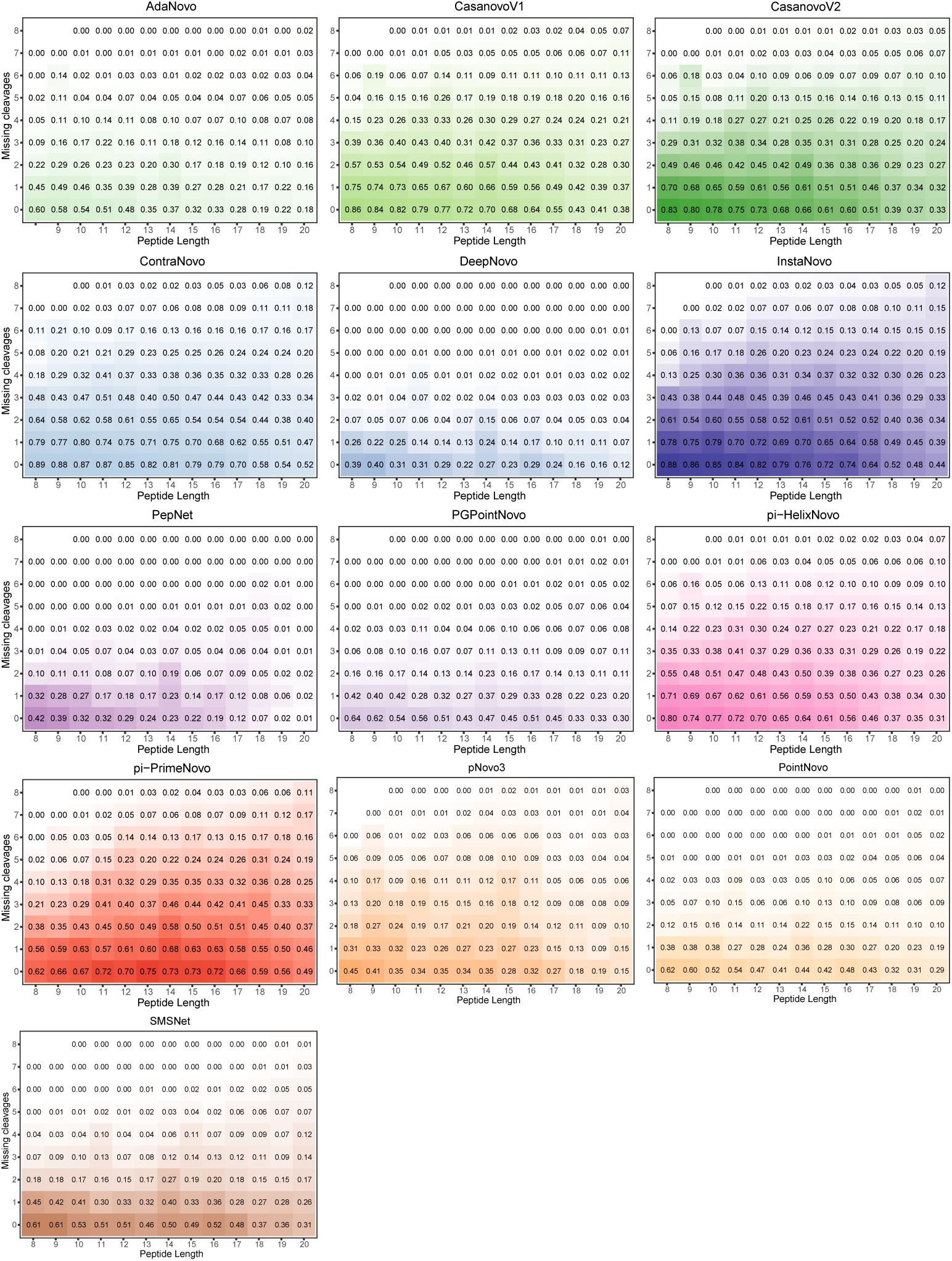
Joint influence of peptide length and missed cleavages on peptide recall. Heatmaps illustrate the peptide recall performance of each de novo peptide sequencing algorithm under varying peptide lengths (x-axis) and numbers of missed cleavage sites (y-axis). Across most models, recall decreases with increasing peptide length and missed cleavages, highlighting the compounded difficulty posed by longer and incompletely digested peptides.

**Fig. S4.**
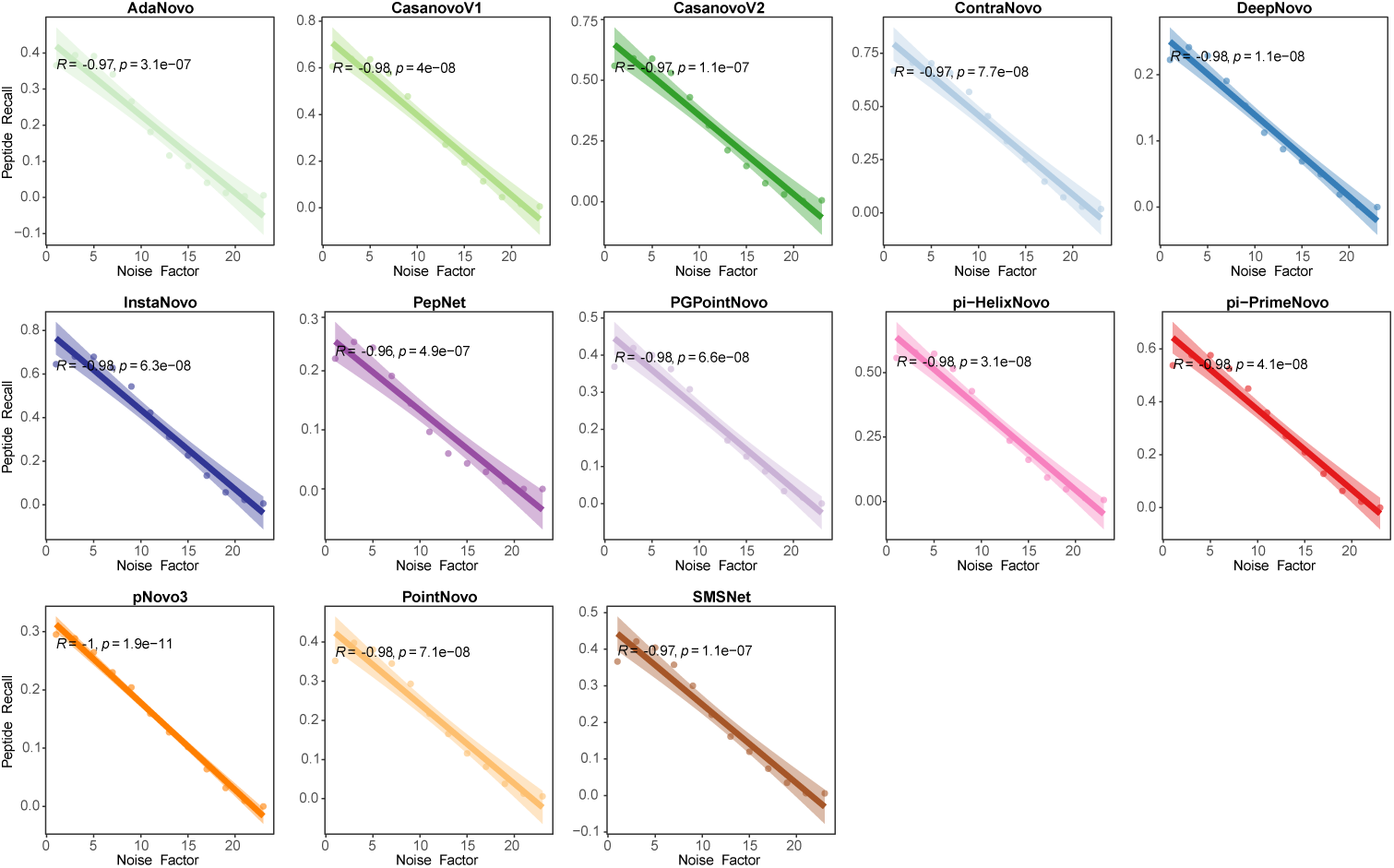
Impact of spectral noise on peptide recall across 13 de novo peptide sequencing tools. Peptide recall declines with increasing noise across all 13 models. Lines show linear fits with 95% confidence intervals.

**Fig. S5.**
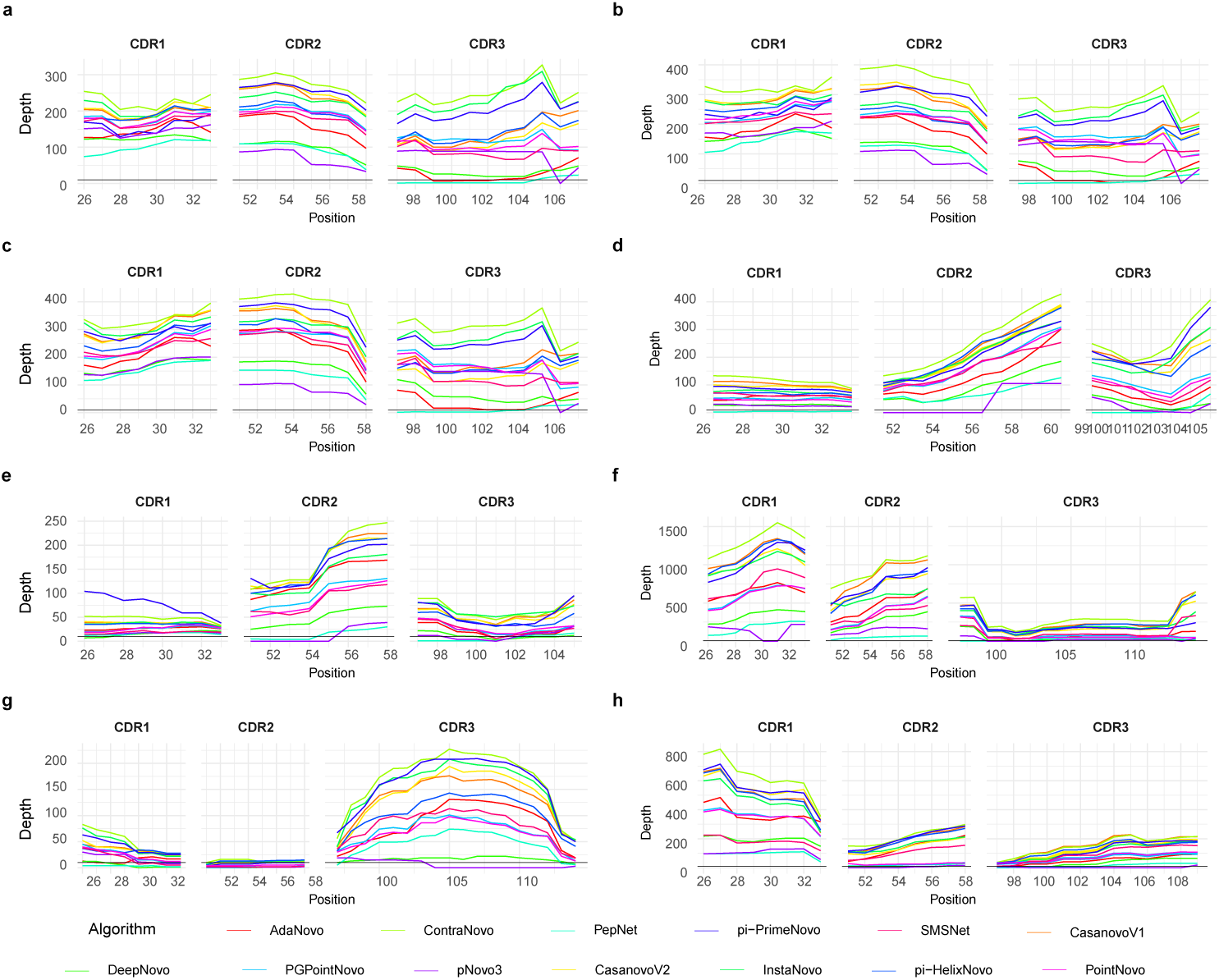
CDR coverage depth for eight monoclonal antibodies across different de novo peptide sequencing algorithms. a–h. Coverage depth of CDR1, CDR2, and CDR3 regions for eight monoclonal antibodies (mAb1–mAb8) across thirteen algorithms. Each line represents a different algorithm, with the depth of coverage shown along the x-axis for peptide positions and the y-axis for the corresponding depth. Algorithms are color-coded as indicated in the legend.

**Fig. S6.**
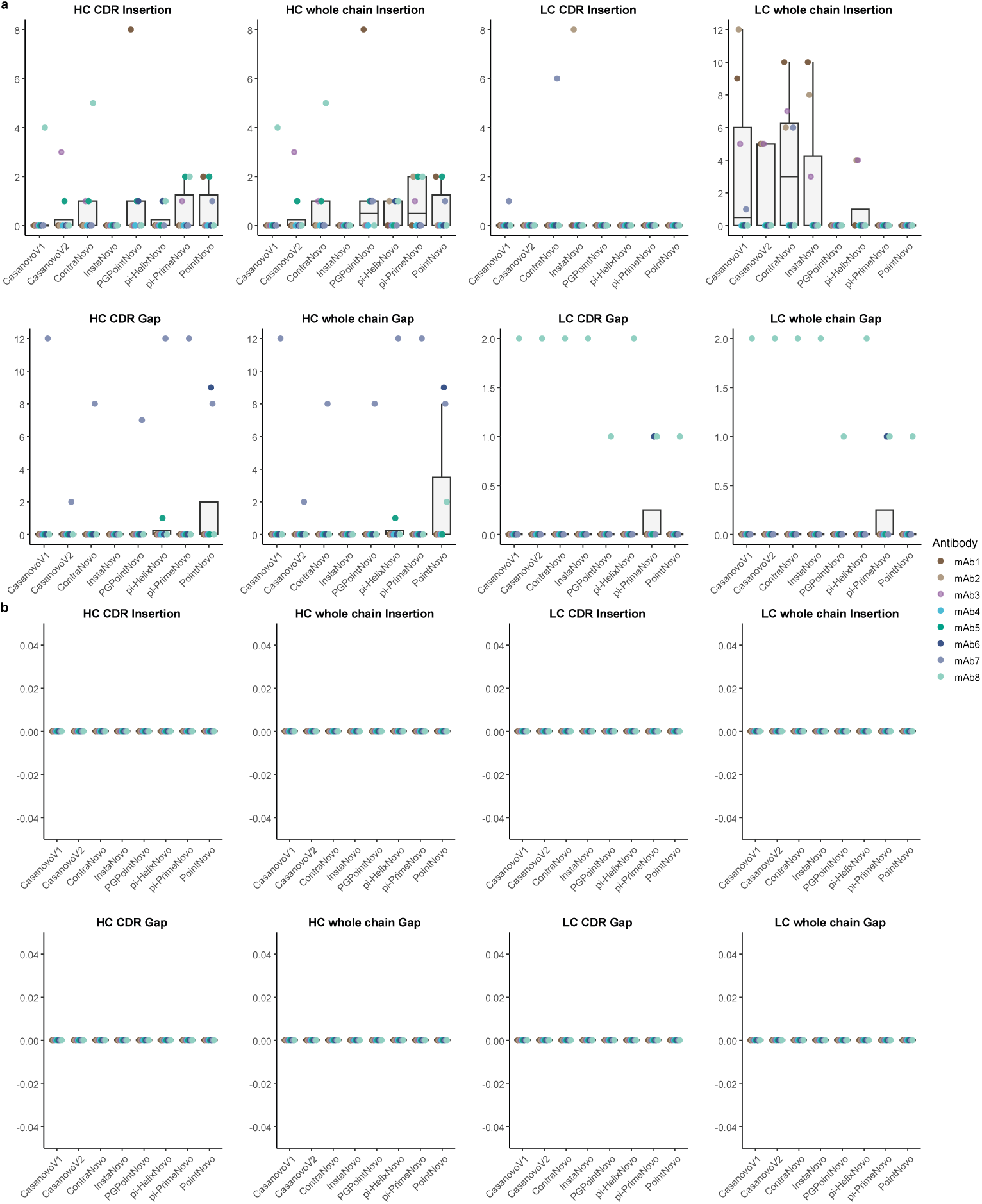
Insertion and gap analysis for Stitch and Fusion. Insertion and gap counts of HC and LC CDRs, and full-length chains, across eight monoclonal antibodies (mAb1–mAb8), utilizing the **a)** Stitch and **b)** Fusion assembly methods.

**Fig. S7.**
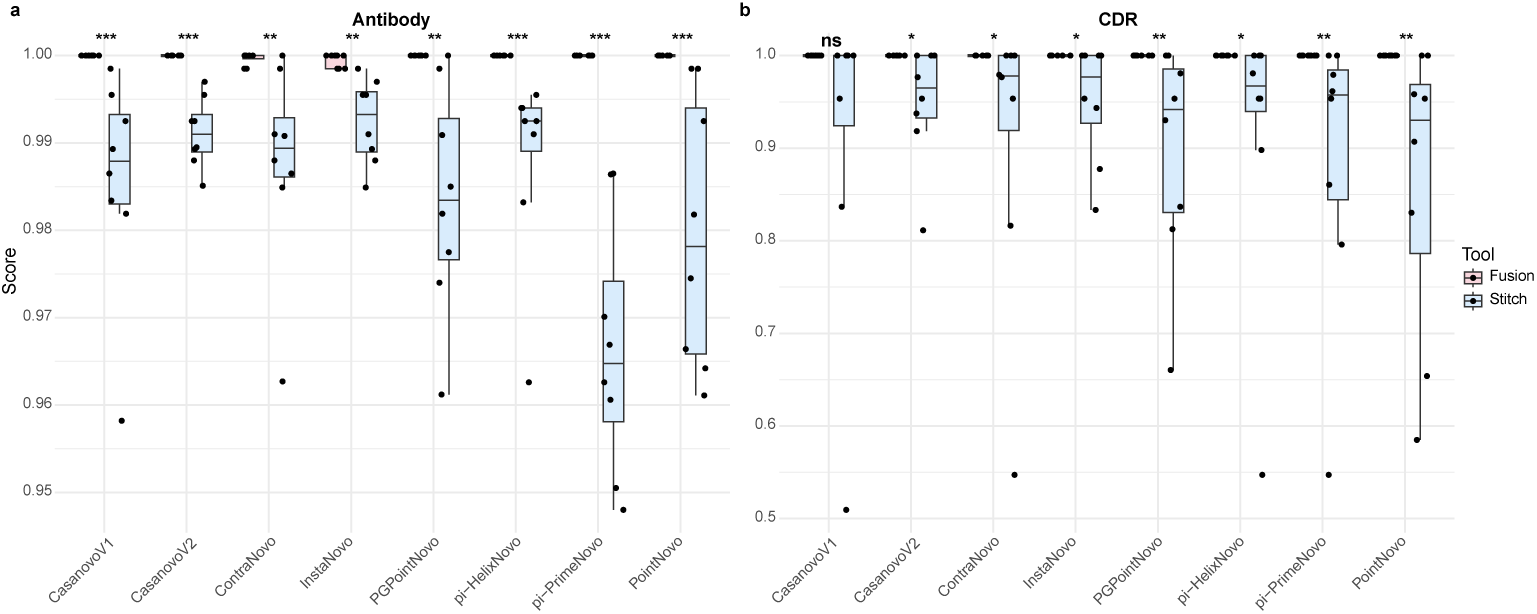
Assembly scores for Fusion and Stitch methods based on predicted sequences from different peptide sequencing algorithms. Assembly scores for **a)** entire antibody sequences and **b)** CDR regions, based on predicted sequences from eight de novo peptide sequencing methods and their assembly results using Fusion and Stitch methods. Wilcoxon test statistical significance is indicated as follows: \**p <* 0.05, \*\**p <* 0.01, \*\*\**p <* 0.001.

**Fig. S8.**
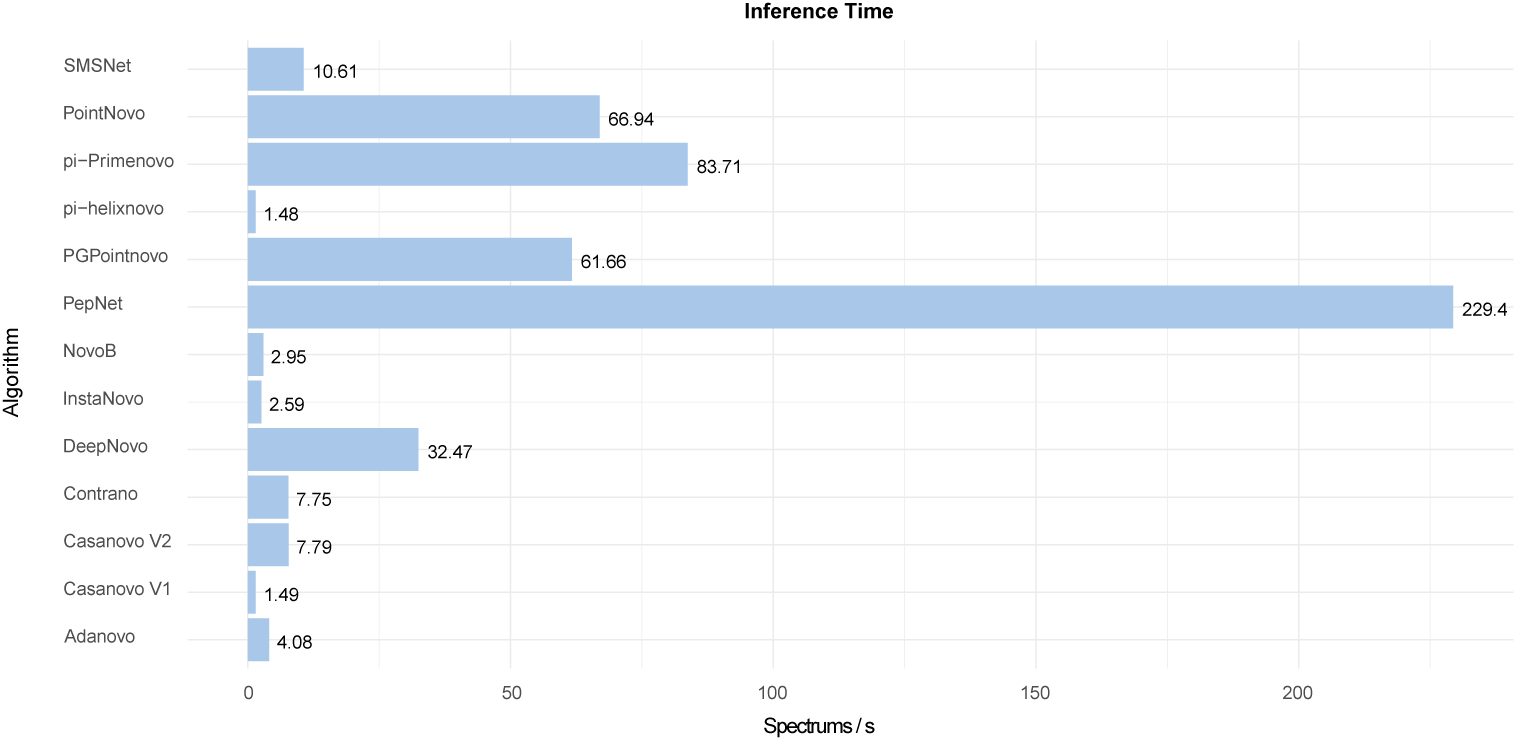
**Inference time (spectra per second) for different de novo peptide sequencing algorithms.**

**Fig. S9.**
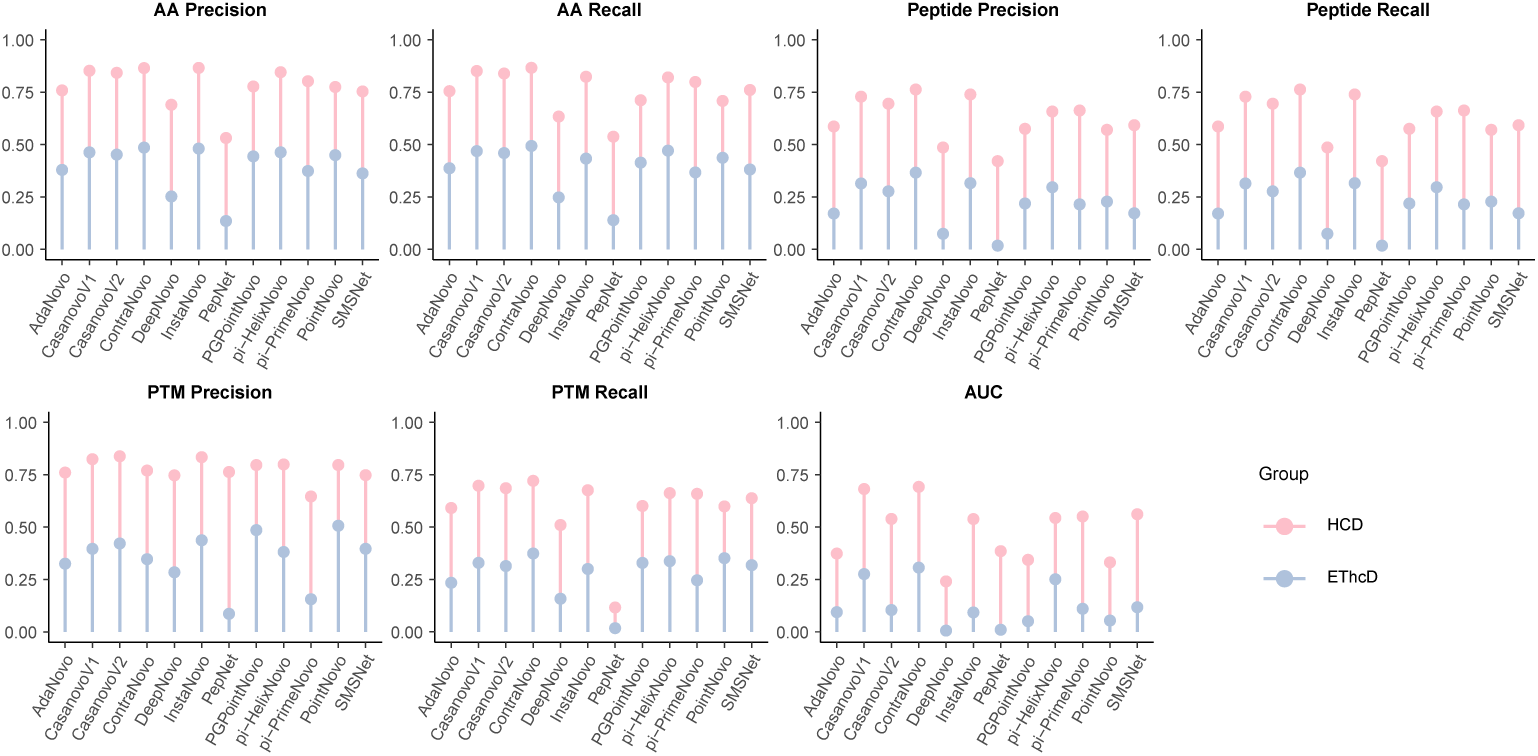
Comparison of accuracy metrics between HCD and EThcD fragmentation modes. Evaluation of different de novo peptide sequencing algorithm performance for AA Precision, AA Recall, Peptide Precision, Peptide Recall, PTM Precision, PTM Recall, and AUC across HCD (pink) and EThcD (blue) fragmentation modes.

**Table S1.** The sequence information of eight assembly mAbs.

| Antibody | LC (AA) | HC (AA) | Species | Protease used |
| --- | --- | --- | --- | --- |
| mAb1 | 217 | 447 | Homo | Asp-N, Chymotrypsin, Elastase, Pepsin, Trypsin |
| mAb2 | 217 | 447 | Homo | Asp-N, Chymotrypsin, Elastase, Pepsin, Trypsin |
| mAb3 | 217 | 447 | Homo | Asp-N, Chymotrypsin, Elastase, Pepsin, Trypsin |
| mAb4 | 214 | 440 | Mus | Asp-N, Chymotrypsin, Elastase, Pepsin, Trypsin |
| mAb5 | 214 | 446 | Mus + Homo | Asp-N, Chymotrypsin, Elastase, Pepsin, Trypsin |
| mAb6 | 214 | 455 | Homo | Asp-N, Chymotrypsin, Elastase, Pepsin, Trypsin |
| mAb7 | 215 | 455 | Homo | Asp-N, Chymotrypsin, Elastase, Pepsin, Trypsin |
| mAb8 | 216 | 450 | Homo | Asp-N, Chymotrypsin, Elastase, Pepsin, Trypsin |

**Table S2.** Overview of all de novo peptide sequencing tools used in this study. For each algorithm, the name of the tool, the model architecture, the year of publication, the project website of the corresponding method, and training status in MassIVE-KB-v1 are displayed.

| Software | Architecture | Year | Ref. | Project website | Retraining status |
| --- | --- | --- | --- | --- | --- |
| DeepNovo | CNN+LSTM | 2017 |  | github.com/nh2tran/DeepNovo | ✓ |
| SMSNet | CNN | 2019 |  | github.com/cmb-chula/SMSNet | ✓ |
| pNovo3 | Post-processor | 2019 |  | http://pfind.net/software/pNovo | — |
| PointNovo | T Net | 2021 |  | github.com/irleader/PointNovo | ✓ |
| Casanovo v1 | Transformer | 2022 |  | github.com/Noble-Lab/casanovo | ✓ |
| PGPointNovo | Other | 2023 |  | github.com/shallFun4Learning/PGPointNovo | ✓ |
| InstaNovo | Transformer | 2023 |  | github.com/instadeepai/InstaNovo | ✓ |
| GraphNovo | GNN | 2023 |  | github.com/AmadeusloveIris/GraphNovo | × |
| PepNet | CNN | 2023 |  | github.com/lkyltal/PepNet | ✓ |
| π-HelixNovo | Transformer | 2024 | | github.com/PHOENIXcenter/pi-HelixNovo | ✓ |
| ContraNovo | Transformer+CL | 2024 |  | github.com/BEAM-Labs/ContraNovo | × |
| NovoB | Transformer | 2024 |  | github.com/ProteomeTeam/NovoB | × |
| AdaNovo | Transformer | 2024 |  | github.com/Westlake-OmicsAI/adanovo_v1 | ✓ |
| PowerNovo | Transformer + BERT | 2024 |  | github.com/protdb/PowerNovo | × |
| Casanovo v2 | Transformer | 2024 |  | github.com/Noble-Lab/casanovo | ✓ |
| π-PrimeNovo | non-autoregressive Transformer | 2025 | | github.com/PHOENIXcenter/pi-PrimeNovo | × |

**Table S3.** Summary of error types across 13 de novo peptide sequencing algorithms.

| | AdaNovo | CasanovoV1 | CasanovoV2 | ContraNovo | DeepNovo | InstaNovo | PepNet | PGPointNovo | π-HelixN |
| --- | --- | --- | --- | --- | --- | --- | --- | --- | --- |
| Number of total predictions | 1,634,030 | 1,638,197 | 1,633,380 | 1,638,248 | 1,589,588 | 1,547,951 | 1,638,248 | 1,589,468 | 1,638,248 |
| Number of total errors | 1,067,403 | 691,721 | 762,611 | 589,821 | 1,256,648 | 531,397 | 1,308,812 | 1,005,272 | 780,92 |
| Inversion first 3 AAs (%) | 7.16 | 7.46 | 7.87 | 6.89 | 4.55 | 8.10 | 3.72 | 8.47 | 6.30 |
| Inversion last 3 AAs (%) | 7.19 | 5.72 | 6.56 | 5.61 | 5.59 | 4.90 | 4.06 | 7.06 | 5.11 |
| Inversion first and last 3 AAs (%) | 0.58 | 0.33 | 0.41 | 0.35 | 0.50 | 0.36 | 0.24 | 0.63 | 0.51 |
| 1 AA replaced by 1 AA or 2 AAs (%) | 2.66 | 3.85 | 3.99 | 5.34 | 0.97 | 7.20 | 4.11 | 2.08 | 3.89 |
| 2 AAs replaced by 2 AAs (%) | 5.16 | 6.02 | 6.25 | 6.52 | 3.04 | 7.38 | 6.81 | 6.12 | 5.51 |
| 3 AAs replaced by 3 AAs (%) | 3.76 | 3.72 | 3.96 | 3.68 | 3.13 | 3.92 | 7.16 | 4.00 | 3.57 |
| 4 AAs replaced by 4 AAs (%) | 3.43 | 3.16 | 3.28 | 3.02 | 3.73 | 3.09 | 5.49 | 3.64 | 3.07 |
| 5 AAs replaced by 5 AAs (%) | 3.30 | 2.80 | 2.85 | 2.67 | 3.57 | 2.60 | 4.57 | 3.55 | 2.62 |
| 6 AAs replaced by 6 AAs (%) | 3.17 | 2.96 | 2.82 | 2.90 | 3.28 | 2.37 | 3.72 | 3.25 | 2.67 |
| More than 6 AAs wrong (%) | 39.37 | 37.69 | 36.18 | 34.57 | 57.72 | 31.24 | 42.83 | 40.60 | 38.81 |
| Other (%) | 24.22 | 26.30 | 25.84 | 28.46 | 13.92 | 28.84 | 17.31 | 20.59 | 27.95 |

## 3 Discussion

In this study, we introduce AbNovoBench, the first comprehensive and standardized benchmarking framework specifically designed for de novo sequencing of mAbs. This framework is built upon a high-quality dataset generated through our previously developed SP-MEGD method[32]. The antibody-specific dataset comprises two essential components: a collection of over 1.6 million high-quality PSMs derived from 131 antibodies across six species, processed with 11 different proteases; and a separate set featuring eight fully annotated antibodies with known sequences and complete MS/MS data for downstream sequence reconstruction. Leveraging this rich data resource, AbNovoBench incorporates a unified training protocol and comprehensive evaluation methodology that enables rigorous and controlled comparisons across a wide array of peptide sequencing algorithms and assembly strategies. This encompasses the entire workflow from spectral interpretation to full-chain sequence reconstruction. By providing a standardized platform, AbNovoBench serves as an invaluable tool not only for fostering continuous improvements in peptide sequencing accuracy and assembly completeness but also for promoting advancements within the field of monoclonal antibody research.

In our analysis of de novo peptide sequencing for antibodies, transformer-based architectures have emerged as the dominant solution. ContraNovo, Casanovo v1/v2, and InstaNovo consistently outperformed recurrent neural network- and graph-based models across a range of accuracy metrics, including precision and recall at the amino acid, peptide, and PTM levels, as well as the area under the precision–recall curve (AUC). Notably, while ContraNovo was trained on the comprehensive 30-million-PSM MassIVE-KB-v1 dataset encompassing a wide range of PTMs, Casanovo v1—trained on a subset comprising approximately 2 million PSMs with only three antibody-relevant PTMs—demonstrated comparable or superior PTM precision to that of ContraNovo. This contrast underscores the value of domain-informed constraint: limiting the modification space reduces overfitting while preserving overall accuracy. Although the training data exclusively comprised tryptic peptides, our analysis demonstrated that DL tools can be effectively applied to various enzymatic datasets, albeit with varying degrees of performance reduction. Importantly, this variation does not adversely affect downstream sequence assembly tasks. However, the presence of missing fragmentation sites, noisy spectra, and long length peptide sequences significantly impairs the performance of all de novo peptide sequencing tools. Notably, when the peptide length reaches or exceeds 25 amino acids, all models demonstrate an almost complete inability to effectively resolve such cases. Additionally, coverage depth analysis indicated that the variable regions of antibodies, especially the heavy chain CDRs, have lower coverage depths than the constant regions. This implies that current peptide sequencing algorithms primarily focus on conserved domain sequences. Therefore, there is a persistent demand for de novo peptide sequencing algorithms specifically designed for antibodies.

Despite ongoing efforts and advancements in de novo peptide sequencing, the reliable assembly of antibodies has long been regarded as a challenging endeavor. However, our findings demonstrate that the template-guided approach (Fusion[29]) achieved error-free reconstruction when supplied with sufficiently dense and high-quality peptide predictions from Casanovo v1/v2, PGPointNovo, PointNovo, *π*-HelixNovo, and *π*-PrimeNovo. Among these sequencing methods, PGPointNovo and PointNovo strike an optimal balance between high fidelity and favorable runtime, rendering them particularly appealing for large-scale applications. During assembly, the coverage depth—particularly within the CDR regions—quantified using a 7-mer sliding window, emerged as the most significant predictor of reconstruction fidelity. Although Fusion assembler can effectively retrieve complete monoclonal antibody sequences from model organisms, fully de novo assembly (ALPS[32]) frequently yields fragmented or ambiguous contigs. Ongoing advancements in experimental techniques and peptide sequencing algorithms may facilitate the realization of fully de novo assembly for non-model organisms.

Nevertheless, several limitations warrant consideration. First, because the benchmark is built almost entirely on Orbitrap Eclipse HCD data collected in data-dependent acquisition mode, our preliminary test on a Trypsin EThcD data set revealed a marked drop in amino-acid and peptide-level accuracy across all tools (Fig. S9), suggesting that performance may diverge under alternative fragmentation or acquisition strategies and motivating future expansion to EThcD, UVPD, and DIA workflows. Second, Fusion and Stitch resolve the isoleucine/leucine (I/L) ambiguity with different evidence streams: Fusion[29] integrates w-ion signals, protease-cleavage specificity, and template context, whereas Stitch[33] infers w-ions from its EThcD-based peptide-sequencing results and then combines these ions with template alignment. Because de novo peptide sequencing accuracy on EThcD spectra is still suboptimal, we normalised both reference and predicted sequences by converting every I residue to L, thereby avoiding an artificial disadvantage to Stitch yet precluding a direct assessment of each assembler’s intrinsic I/L-discrimination capability. Future versions of AbNovoBench will revisit this issue as improved de novo algorithms for alternative fragmentation modes become available and as broader training data sets are incorporated.

In conclusion, AbNovoBench fills a long-standing void in antibody de novo sequencing research by providing a scalable, antibody-centric benchmarking platform. Transformer architectures currently define the state of the art, and Fusion stands out as the assembler of choice for high-fidelity antibody reconstruction. Future work should broaden benchmark coverage to alternative mass-spectrometry modalities, investigate transfer-learning strategies across instrument types, and foster community-driven updates via the AbNovoBench portal, ensuring that de novo sequencing tools evolve in tandem with advances in antibody biology and proteomic technologies.

## 4 Methods

### 4.1 Ethical approval declarations

Animal experiments were carried out in accordance with the approval of the Institutional Animal Care and Use Committee at Xiamen University (XMULAC20230310) and by the Guide for the Care and Use of Laboratory Animals.

### 4.2 Monoclonal antibody sample preparation

The mAbs used for this study were expressed by ExpiCHO-S^™^ suspension expression system (Thermo Fisher Cat#A29133) and subsequently purified via Protein A column (Cytiva). Next, we employed the single-pot multi-enzymatic gradient digestion (SP-MEGD) method, developed in our previously study [32], for mAb sample preparation. SP-MEGD enables efficient “head-to-tail” peptide assembly and requires as little as 50 µg of sample, making it suitable for routine analysis. The proteases utilized in this study include Asp-N, Chymotrypsin, Elastase, Pepsin, Trypsin, Glu-C, aLP, Lys-C, Thermolysin, as well as combinations of Trypsin with Asp-N and Trypsin with Glu-C. Initially, the lysis buffer [6 M guanidine hydrochloride (Gu.HCl), 20 mM dithiothreitol (DTT), 100 mM Tris, pH 8.5] was added to 200 µg of purified mAb at an approximate ratio of 1:1 (microliters of lysis buffer to micrograms of protein). The mixture was denatured and reduced at 60*^◦^*C for 30 min followed by alkylation with 40 mM iodoacetamide (IAA) in the dark for 30 min at room temperature. The alkylated antibody sample was transferred into a solution containing 0.8 M Urea, 50 mM Tris-HCl, pH 8.0 in a 10k filter (Millipore, USA) at 4*^◦^*C to eliminate interfering substances that might influence enzymatic digestion. The proteolytic enzymes, namely trypsin, chymotrypsin, pepsin, elastase, and Asp-N, were added at a ratio of 1:20 (w/w). The digestion solution was incubated at 37*^◦^*C for 6h and samples were collected at two-hour intervals respectively. After digestion, the reaction mixture was quenched with 10% trichloroacetic acid (TFA) for 30 min at 37*^◦^*C. The supernatant was desalted by Sep-Pak C18 Vac cartridges (Waters) following the instructions before being lyophilized to dryness or stored at −20*^◦^*C or dissolved in 0.1% FA for LC–MS/MS analysis.

### 4.3 LC–MS/MS analysis

The digested peptides were separated through reversed-phase chromatography on a Vanquish^™^ Neo UHPLC (column packed with PepMap^™^ 100 C18, 75 µm*×*50 cm, 2 µm, Thermo Fisher Scientific, USA) coupled to an Orbitrap Eclipse mass spectrometer. Samples were eluted over a 70-min gradient from 0 to 35% solvent B (0.1% formic acid in 80% acetonitrile) at a flow rate of 300 nL/min. Solvent A was 0.1% formic acid in water. Full MS1 scans were acquired over a range of m/z 350–2000 with a resolution of 120,000. MS1 scans were obtained with a standard automatic gain control (AGC) target and a maximum injection time of 100 ms. The precursors were fragmented by stepped high-energy collision dissociation (HCD). The stepped HCD fragmentation included steps of 27%, 35%, and 40% normalized collision energies (NCE). MS2 scans were acquired at a 30,000 resolution, a 5E4, AGC target, and a 250 ms maximum injection time. In total, 141 bottom-up mass spectrometry datasets were generated for this study.

### 4.4 Preprocessing

The raw files need to be converted to open-format files to be compatible with de novo sequencing tools[34]. We reformatted the raw MS/MS data files from the previously mentioned data sets to Mascot Generic Format (MGF) using ProteoWizard[35]. A MGF file stores the m/z and intensity pairs of multiple mass spectra in a single text format. De novo peptide sequencing tools predict amino acid residues by accessing the mass differences between the MS/MS peaks[36].

### 4.5 Parameters for de novo peptide sequencing algorithms

To perform a fair comparison between spectrum-graph-based tools like pNovo3[33], which are released only with pre-trained models, and the DL algorithms, we trained all DL-based tools on high-resolution MS/MS data from the human proteome using the HCD library from MassIVE KB v1[37], which consists of 1,913,865 peptide-spectrum match (PSM) from 1,377,087 different peptides. The filtered dataset was downloaded from the official MassIVE repository (https://massive.ucsd.edu/ProteoSAFe/DownloadResult?view=download filtered mgf library&task=3cac03860ff7453a821332ab4cff20f4&show=true). After filtering PSMs with unrelated variable modifications, we split the spectra into training, and validation sets at a ratio of 95:5 while making sure that the split data sets did not share any common peptides. Each model was trained for 20 epochs using pre-defined parameters from each tool. We executed all tools at a precursor tolerance of 10 ppm and fragment mass tolerance of 0.02 Da. For each algorithm, carbamidomethylation of cysteine (C+57.02 Da) was set as a fixed modification. Oxidation of methionine (M+15.99 Da) and deamidation of asparagine and glutamine (N+0.98 Da and Q+0.98 Da) were set as variable modifications. All deep learning–based models were trained on a Linux server equipped with 128 CPU cores, 252 GB of RAM, and a single NVIDIA RTX 3090 GPU, except for PrimeNovo and GraphNovo, which were trained on a high-performance computing node comprising 108 CPU cores, 840 GB of RAM, and eight NVIDIA A100 GPUs (40 GB each). During the inference stage of de novo peptide sequencing, all models were executed on the same Linux server with 128 CPU cores, 252 GB of RAM, and a single NVIDIA RTX 3090 GPU. We executed pNovo3[33] on a Windows 64-bit computer since the software was not supported by a Linux operating system.

### 4.6 Assembly of identified peptides

To investigate whether peptides predicted by different DL algorithms can be assembled into a full length HC or LC antibody sequence without prior knowledge of ground truth sequences, we used the software Stitch v1.5[31] and Fusion[32] (XA-Novo platform, https://xa-novo.com/), both of which are template-based sequence assemblers for antibody peptides. Since each de novo peptide sequencing algorithm computes its confidence score using distinct methodologies, we established the threshold for the confidence score based on amino acid-level precision. All de novo peptides generated by the DL algorithms, with prediction scores exceeding the minimal thresholds that ensure cumulative amino acid-level precision of at least 50% and 90%, were utilized for Fusion and Stitch, respectively. The germline antibody sequences from IMGT are employed as templates to guide the assembly of de novo peptide reads.

### 4.7 Evaluation metrics

For validating de novo peptide sequencing algorithms, we compared each prediction to a pseudo-ground truth, which is commonly obtained by database search[36, 38]. Since the evaluated data sets lack a labeled ground truth for each spectrum, we generated labels for each spectrum using PEAKS AB[29]. To measure the accuracy of a given model’s prediction, we compared the real peptide sequence and the de novo peptide sequence of each spectrum at both the amino acid and peptide levels. At the amino acid level, a prediction is considered correct if its mass differs by less than 0.1 Da from the ground truth and if its prefix or suffix differs by no more than 0.5 Da. Precision (Eq. (1)) and recall (Eq. (2)) are then calculated as the ratio of correctly predicted amino acids to the total predicted and ground truth amino acids, respectively. At the peptide level, a prediction is correct only if all amino acids in the peptide match the ground truth. Precision (Eq. (3)) and recall (Eq. (4)) are then calculated as the ratio of correctly predicted peptides to the total predicted and ground truth peptides, respectively. Similar to amino acid-level metrics, PTMs identification precision (Eq. (5)) and recall (Eq. (6)) can be formulated as the ratio of correctly predicted PTMs to the total predicted and ground truth PTMs, respectively. Most importantly, all sequencing tools report confidence scores for their predictions. The confidence scores reflect the quality of predicted amino acids and are valuable for downstream analysis [e.g., reconstructing the entire protein sequence from its peptides [28720701]. Setting a higher threshold of confidence scores will output a smaller set of peptides with high precision but will leave the rest of the dataset without results, hence leading to lower recall and vice versa. Hence, given the availability of recall, precision, and confidence scores, it is reasonable to draw precision-recall curves and use the area under the curve (AUC) as a summary of de novo peptide sequencing accuracy.

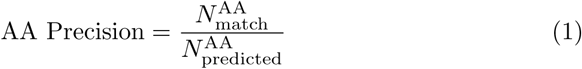

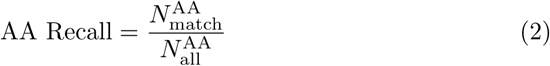

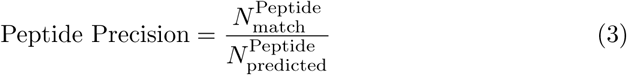

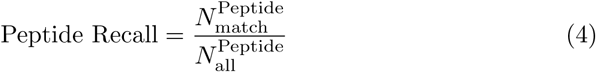

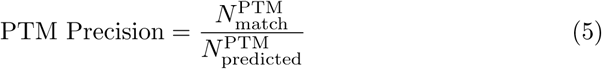

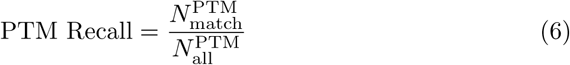

To evaluate the amount of noise and missing fragment ions in each spectrum, we labeled each peak as a peptide peak or a noise peak using the Pyteomics framework[39].

For each cleavage site, we tried to identify twelve different ion types (a, a(2+), a-NH_3_, a-H_2_O, b, b(2+), b-NH_3_, b-H_2_O, y, y(2+), y-NH_3_ and y-H_2_O) since all evaluated de novo peptide sequencing algorithms take these ion types into consideration for retrieving the peptide sequence. If possible, we matched these ion types to corresponding peaks in the spectra within a tolerance of 0.5 Da. Otherwise, we declared the cleavage site as missing. We only considered noise peaks if their intensity exceeded the median noise intensity for each data set. The number of noise peaks above this threshold was used to calculate the noise factor, which is defined as the ratio of the number of high-intensity peaks and the number of fragment ion peaks.

Additionally, we assess the models’ potential for assembly, specifically focusing on accurate coverage depth in relation to the determined mAb sequence. It’s an important influencing factor while seldom be considered. The subsequent analysis concentrated on characterizing antibody assembly quality. Specifically, sequence coverage was defined as the percentage of amino acids in the reference antibody sequence that were covered by the assembled contig, while sequence accuracy measured the proportion of correctly annotated residues within the assembled region. To comprehensively evaluate the performance of sequence assembly, we designed a unified scoring formula that is applied independently to two catagories of interest: the whole antibody and the CDR region. For each catagory, the score is computed as a length-weighted average of the chain-level quality (Eq. (7)), incorporating coverage (C), accuracy (A), insertion

I. and deletion (G) errors, as well as the region-specific sequence length (L):

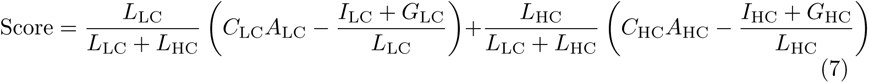

where the subscripts LC and HC denote the light and heavy chains, respectively.

All variables are specific to the selected region—either the full chain or the CDR segment. This dual-region scoring framework enables a comprehensive evaluation of reconstruction quality, capturing both the global assembly performance and the local accuracy in functionally critical regions.

To classify the assembly results generated by the de Bruijn assembler ALPS[29], we aligned the target contigs with the reference antibody sequence using BLAST[40]. For Fusion and Stitch, we employed the Needleman-Wunsch[41] algorithm in conjunction with the BLOSUM60 matrix to align the assembled sequences with their corresponding antibody sequences, achieving the most intuitive results.

## Acknowledgements

This work was supported by the National Science and Technology Major Project for Innovative Drug Research and Development (2025ZD1803703 to Q.Y.); the National Natural Science Foundation of China (32401237 to YX. and 92369110 to Q.Y.); the Fujian Provincial Natural Science Foundation of China (2024J08358 to Y.X.); the Natural Science Foundation of Xiamen, China (3502Z202371039 to Y.X.); the State Key Laboratory of Vaccines for Infectious Diseases, Xiang An Biomedicine Laboratory (2025XAKJ0200001 to R.Y.); and the Scientific Research Foundation of the State Key Laboratory of Vaccines for Infectious Diseases (2024SKLVDzy06 to Y.X.).

